# A First Order Phase Transition Underlies the Formation of Sub-Diffractive Protein Aggregates in Mammalian Cells

**DOI:** 10.1101/148395

**Authors:** Arjun Narayanan, Anatoli B. Meriin, Michael Y. Sherman, Ibrahim I. Cissé

## Abstract

Failure in protein quality control can often lead to protein aggregation, yet in neurodegenerative diseases, by the time aggregates can be seen, the cells have advanced well into the disease pathology. Here, we develop a quantitative imaging approach to study the protein aggregation process in living mammalian cells with unprecedented spatio-temporal resolution. We find that sub-diffractive precursor aggregates may form even in untreated cells, and their size distribution is exactly as predicted for a system undergoing a first order phase transition. Practically, this implies that as soon as aggregates reach a critical size (*R_c_* = 162 ± 4 nm untreated cells), they will spontaneously grow into large inclusions. Our data suggest that a previously uncharacterized, RuvBL1 dependent mechanism clears aggregates above the critical size. Our study unveils the existence of sub-diffractive aggregates in living cells; and the strong agreement between cellular data and a nucleation theory, based on first order phase transition, provides insight into regulatory steps in the early stages of aggregate formation in vivo.

---

Neurodegenerative diseases, such as Parkinson’s Disease, Amyotrophic Lateral Sclerosis, and Alzheimer’s Disease, are characterized by the appearance of large protein aggregates in cells and in the extracellular space (*1*). It is hypothesized that intermediate species in the aggregation process are likely more toxic moieties (*2-9*) than conventionally visible large aggregates, plaques or fibres. However detecting and characterizing intermediate aggregates remains a fantastic technical challenge. Capturing the early steps of protein aggregation in living cells can help uncover hidden mechanisms in their formation and regulation in vivo, as well as elucidate their putative roles in protein misfolding diseases (*2-9*).

Here we develop a quantitative super-resolution assay to study the early steps protein aggregation in mammalian cells. We adopt proteasome inhibition as an approach used to study the formation of large aggregates in living mammalian cells (*10-13*). Treatment of cells with the proteasome inhibitor MG132 (*14*) leads to the gradual accumulation of misfolded, aggregation-prone proteins, and to the formation of the aggresome, a large juxta-nuclear inclusion body akin to Lewy bodies (*15, 16*) in Parkinson’s disease cells. We engineered mammalian cell lines expressing Synphilin 1 - a marker of aggregates in Parkinson’s disease (*10, 17, 18*) - fused to a fluorescent protein Dendra2 (*19*). Dendra2 is a green to red photo-convertible protein that enables photo-activation localization microscopy (PALM) (*20*), a single-molecule based super-resolution (*20-22*) approach we used previously to quantitatively image protein clustering with high spatio-temporal resolutions in living cells (*23, 24*).

Imaging Synphilin 1 by conventional fluorescence, shows emergence of the aggresome around 135 minutes after treatment (Fig. 1A-E, top panels and **supplementary movie 1)** but formation dynamics of the aggresome cannot be readily measured with this imaging approach. Instead, when we perform live cell super-resolution imaging (*25*) (Fig. 1A-E, bottom panels), we can detect and quantify the growth of individual aggresomes (Fig. 1F), from their inception at length scales unattainable in previous live cell studies (*26*). Furthermore, in addition to the large aggresome, the examination of Fig. 1A-E (bottom panels and **supplementary movie 1**) shows a population of sub-diffractive aggregates throughout the cellular cytoplasm and indiscernible in the conventional images (top panels). Thus, our live cell super-resolution imaging approach reveals a previously undetected population of sub-diffractive aggregates.

**Figure 1:**
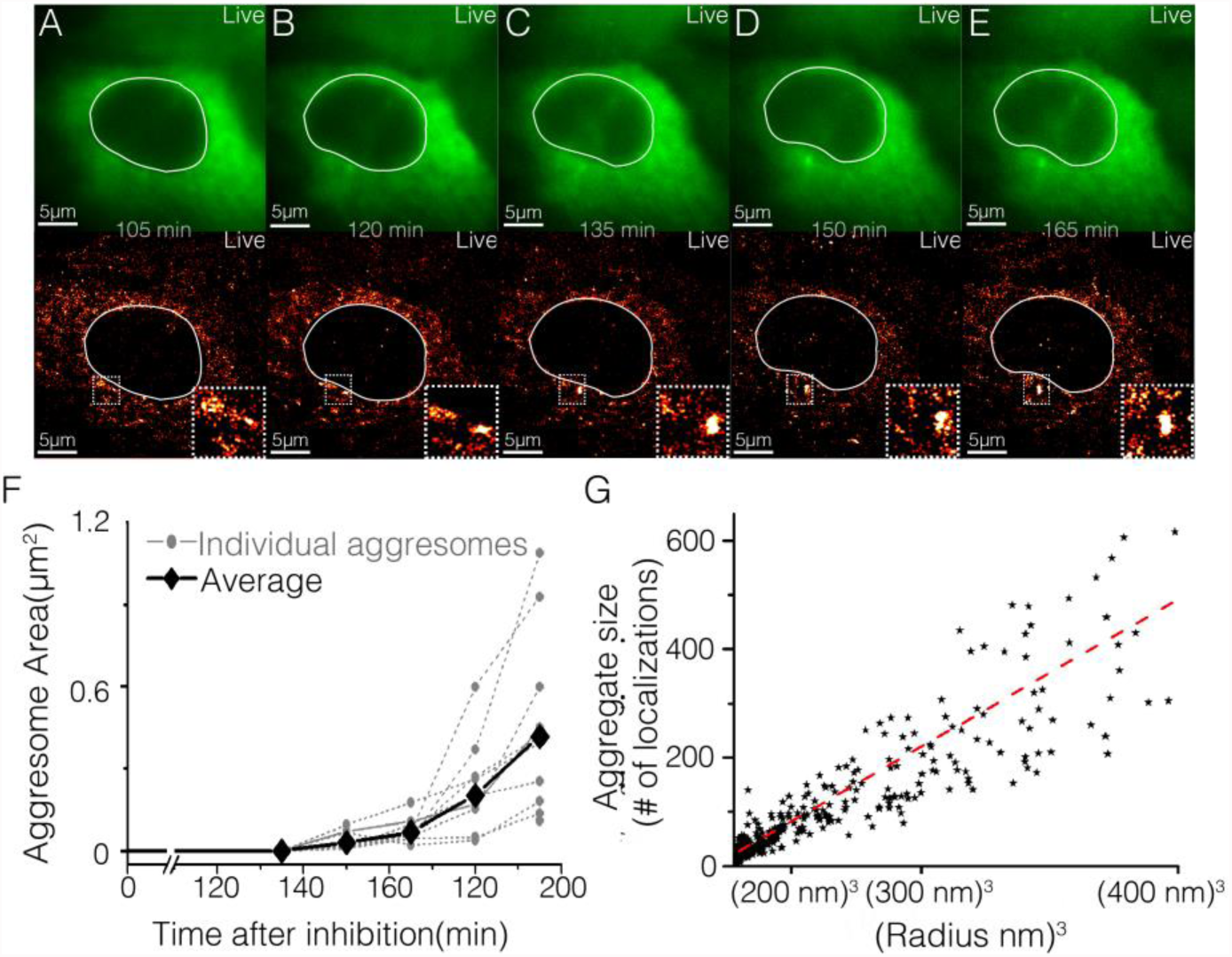
Live cell super-resolution imaging unveils sub-diffractive aggregates in addition to classically observed perinuclear inclusion. ***A-E:*** A single living cell expressing Dendra2-Synphillin as observed by conventional imaging (top panels, green) and time-lapse super-resolution reconstructions (bottom panels, red-hot color code) is represented for select time points between 105 minutes to 165 minutes after treatment with proteasome inhibitor MG132. White line delineates cell nucleus; a red-hot color code is used to represent the local density of detections. Both conventional and super-resolution images show the gradual formation of a large perinuclear inclusion (the aggresome, see insets in bottom panel).. **F:** The time-dependent sizes of individual aggresomes (gray dots) are measured from super-resolution images of ten cells, illustrating cell to cell variability in the growth dynamics; the average time dependent size from the ten cells is also plotted (black diamonds). In addition to the aggresomes, super-resolution images (***A-E*** bottom panels) reveal many sub-diffractive aggregates profusively distributed throughout the cytoplasm, that were not observed by conventional imaging (***A-E*** top panels). **G:** The sub-diffractive aggregate size as measured by number of super-resolution localization events per aggregate is plotted against the spatial size (radius cube) of aggregate. The data suggests that size of sub-diffractive aggregates scales with the volume of the aggregates, suggesting a uniform density of Dendra2-Synphilin detections throughout the aggregates, up to the z-axial (400 nm) cut-off of our super-resolution microscope (see **supplementary text 1** for details). Data in G includes N= 4000 aggregates from 6 cells imaged 120 minutes after MG132 treatment.

We characterize the properties of these sub-diffractive aggregates using density based spatial clustering of applications with noise (DBSCAN) (*27*). We record for each aggregate, the radius, and the number of localization events (fluorescence detection events) ((*25*) and **supplementary text 1**). Only aggregates with a radius greater than our localization accuracy (estimated to be ~20nm (*23*)) are interpreted in our analysis; aggregates of radius less than 25nm are discarded.

As represented in Fig. 1G, we find that the number of localization events per aggregate is proportional to the radius cubed (volume) of the aggregate (also see **supplementary text 1**). This observation implies that sub-diffractive aggregates have a defined density, with the aggregate size scaling linearly with volume. Relying on the precise number of molecular detections to estimate aggregate size can be complicated by single molecule photo-physical variability (*28*). Here, we rely on the existence of a well-defined density to use the spatial extent (radius cubed) of the aggregate as the measure of the size. For subsequent theoretical analyses, we found it practical to define the aggregate size as a reduced numerical parameter ‘*n*’ (see **supplementary text 1**, and **Fig. S1**].

Previous studies, from experiments done in vitro, have invoked nucleation and growth as a potential mechanism underlying aggregate formation (*29*). However, such models imply that aggregation occurs through a first order phase transition into a so-called state of *super-saturation,* characterized by a well-defined nucleation barrier (*30*). The nucleation barrier reflects a critical aggregate size above which spontaneous growth is energetically favoured, and below which aggregate disassembly is favoured. Such a critical aggregate size, if it exists, has been difficult to measure experimentally (*29*), due in part to the challenge of detecting the stochastically formed, transient precursor clusters; and it is unclear even if proven in vitro, whether phase transition formalism may still hold inside the cells where complex biological quality control mechanisms exist. If a first order phase transition is pertinent, there are clear theoretical expectations for the distribution and evolution of aggregate sizes. Therefore, we investigate the mechanism behind sub-diffractive aggregates formation and growth in mammalian cells, initially, by studying the size distribution of the aggregates, and how the distribution evolves with time.

In nucleation and growth a system may be either in a *sub-saturated* state (Fig. 2A), or in a *super-saturated* state (Fig. 2B & C). In the first case the formation and growth of sub-diffractive aggregates is not favoured energetically. In such a *sub-saturated* system, an exponential distribution of aggregate sizes is expected (*30, 31*). For a *sub-saturated* state, the overall distribution of aggregate sizes does not change with time even as individual aggregates may grow or disassemble (**supplementary text 2**, and simulations in Fig. 2D).

**Figure 2:**
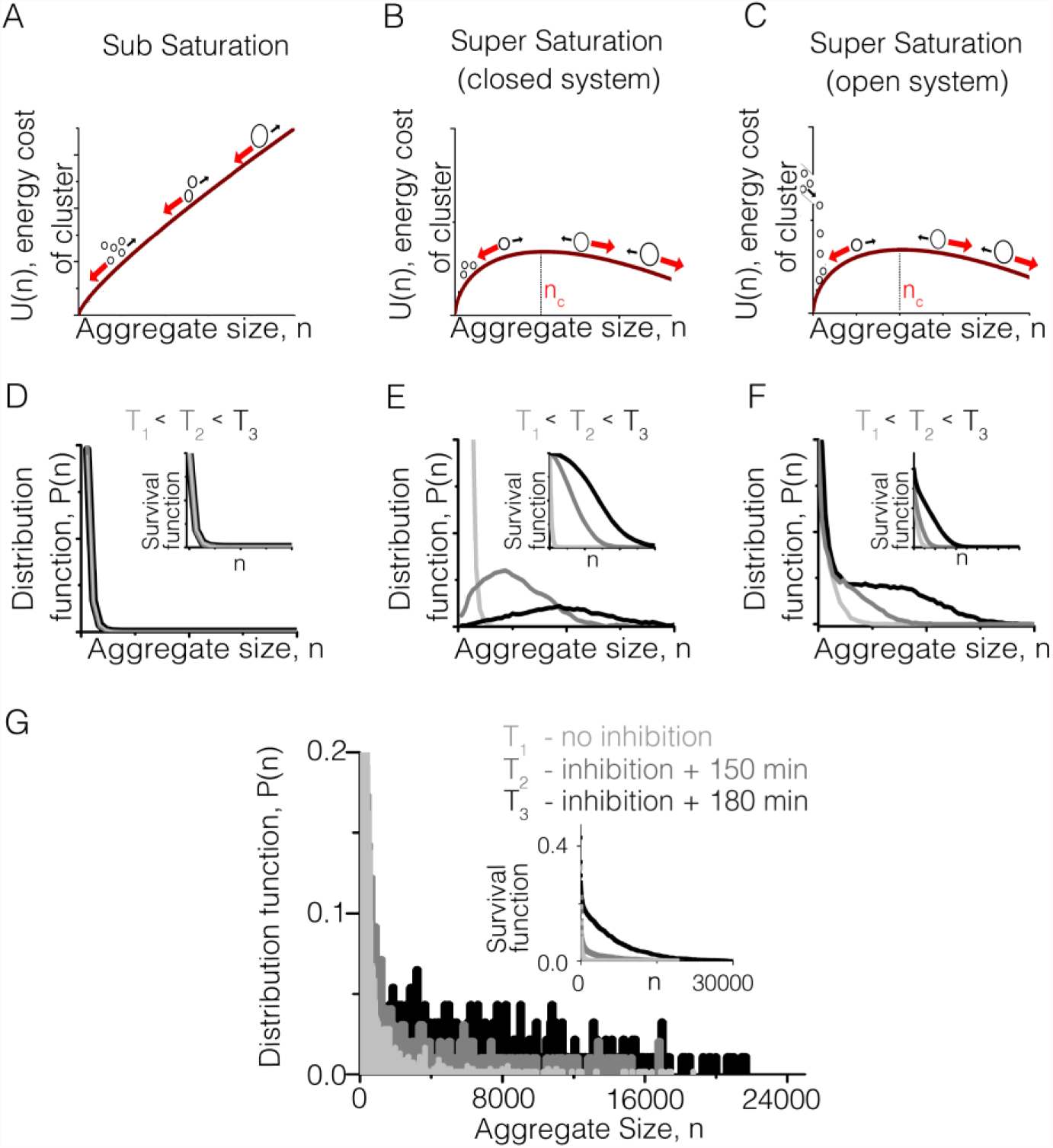
The time evolution of aggregates size distribution is indicative of a super-saturated system: **A-C** Free energy cost of aggregate formation is plotted as a function of aggregate size as predicted by condensation theory in three different regimes. **A:** In a *sub-saturated* system free energy cost increases monotonically with aggregate size, and spontaneous disaggregation is favored (**supplementary text 2**). **B & C:** In a *super-saturated* system, a nucleation barrier exists above which free energy cost decreases with aggregate size, and spontaneous growth is favoured. **D-F:** Simulations of the aggregate size distributions as predicted by the energetics illustrated in **A, B** and **C** respectively. **D** For the *sub-saturated* system (in **A**), an exponential distribution is obtained, independent of time. **E**: For a *super-saturated* system without addition of new constituents (closed system in **B**) a time dependent distribution is obtained where by small aggregates are depleted and the system evolves towards a peak at large aggregate size. **F:** For a *super-saturated* system with addition of constituents, there is no depletion of small aggregate size, but progressive shoulder towards larger aggregate sizes is obtained with time. Insets in **D-F** represents the survival (cumulative distribution) function for the corresponding simulation. **G** Super-resolution data representing the histogram of aggregate sizes from cells fixed and imaged at three representative time points after inhibition by MG132, shows a time evolution consistent with the *super-saturated* system in **C** & **F**. These results indicate that aggregation in the mammalian cells are consistent with *super-saturated* condensing system. Data in **G** represents the normalized histogram from 10000 aggregates (light grey) from 10 untreated cells, 6500 aggregates (grey) from 8 cells and 4000 (dark grey) from 10 cells. Details of simulation in **D-F** in (*25*)

Alternatively, in a *super-saturated* state, the system is poised such that aggregates that stochastically reach the critical size become energetically favoured to grow spontaneously (see **supplementary text 2** and **Fig. S2**). In such a case the distribution of aggregate sizes is time-evolving (simulations in Fig. 2E& F) and may result in a peak at large aggregate sizes when the total pool of contributing proteins is conserved (Fig. 2E). Alternatively, the pool of contributing proteins may be continuously replenished, leading to an exponential distribution of small aggregates coexisting with a growing shoulder at larger aggregate sizes (Fig. 2F); This may likely be the case in living cells when new misfolded proteins can constantly be added to the system. As represent in Fig. 2G our super-resolution data reveals a distribution of aggregate sizes with a shoulder growing towards larger aggregates, as a function of time after treatment, more consistent with the simulations in Fig. 2F. This result suggests that sub-diffractive aggregate formation and time evolution may behave as a *super-saturated* condensation system.

A *super-saturated* system is expected to exhibit precise energetics, underlain by a critical aggregate size (*30*) (noted here as *n_c_* or *R_c_*). *n_c_* (or *R_c_*) is the point at which the surface energy cost is balanced by the minimising energy of the molecules buried in the bulk of the aggregate. In particular the expected form of the free energy cost to form a aggregate of size “n”, is given by *U*(*n*) = *an*^2/3^ − *bn*, with two terms (*an*^2/3^ and *bn*) representing the surface and bulk contributions respectively (**supplementary text 2**, and **Fig. S2**). Moreover, in this formalism, for aggregates below the critical size (i.e *n* < *n_c_*) the Boltzmann distribution (*P*(*n*) ∝ *e*^−*U*(*n*)^), the equilibrium thermodynamics exponentially-suppressed distribution, would be expected even though the full system may not be in equilibrium (*30*).

We examine whether the sub-diffractive aggregates in the mammalian cells truly exhibit such stringent energetics. Given that for a *super-saturated* system the sub-critical aggregate (*n* < *n_c_*) size distribution may be approximated as *P*(*n*) ∝ *e*^−*U*(*n*)^ (*n* < *n_c_*), then the negative logarithm should give the free energy cost *U*(*n*), i.e −*Log*(*P*(*n*)) ∝ *U*(*n*), for *n* < *n_c_*. By plotting the negative logarithm of the size distribution one can test how well the distribution is governed by the precise free energy cost *U*(*n*) = *an*^2/3^ − *bn* (see prediction in Fig. 3A). We find a remarkable agreement between the experimentally measured sub-critical size distribution shown in Fig. 3B and this very specific prediction of simple condensation theory (see (*25*) and **supplementary text 2**).

**Figure 3:**
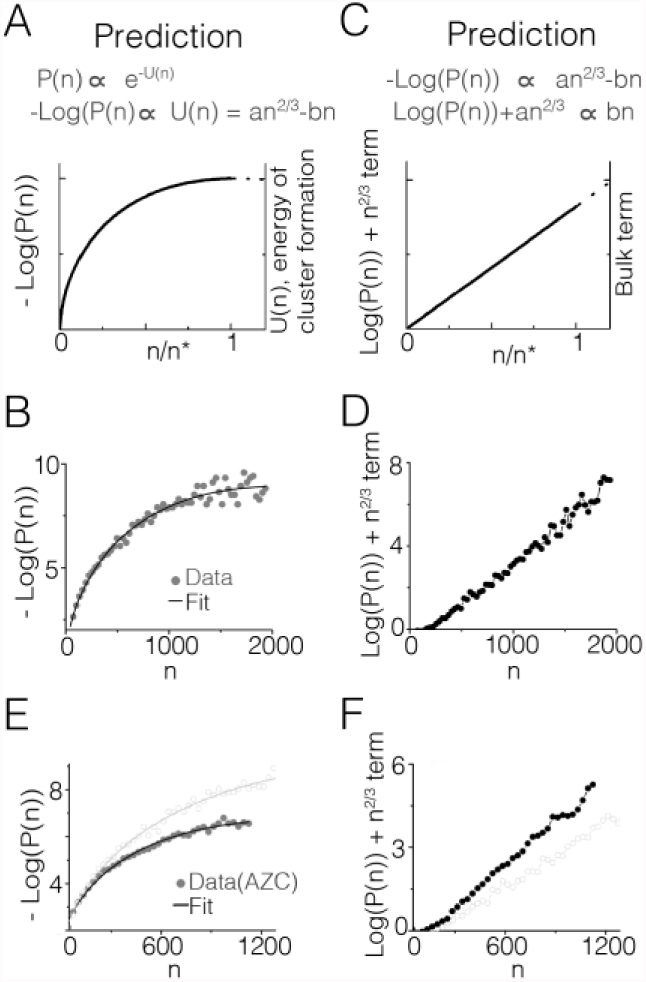
Figure 3: Aggregate size distributions are precisely described by the kinetics of first order phase transition. **A:** The energetics of a *super-saturated* system as expected from theory on the kinetics of first order phase transition [**supplementary text 2**]. Since the expected distribution of subcritical clusters is the boltzmann (exponential) distribution, i.e. *P*(*n*) ∝ *e*^−*U*(*n*)^; *with U*(*n*) = *an*^2/3^ − *bn*, the energy cost dependence on cluster size can be derived from the logarithm of the distribution, i.e. −*Log*(*P*(*n*)). In the energetics, *an*^2/3^ is a measure of surface tension of the condensing aggregates while −*bn* is a measure of the bulk energy gain from n molecules joining an aggregate. **B** Plotting the logarithm of the distribution (histogram) of measured sub-critical aggregates (gray dots) shows the predicted curved dependence with the aggregate size (n), and fits well to the predicted energy form *U*(*n*) = *an*^2/3^ − *bn*. **C**: Theoretically, the surface dominates for the smallest cluster size (i.e. very low n), and subtracting the surface term from the data should result in a predicted linear dependence on cluster size (representing the resultant bulk term) i.e. 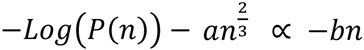 or as represented here 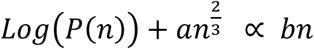. **D**: Subtracting the surface term fit for low n (here n<200) yields a resultant data set that is remarkably linear, further indicating that the specific energetics of first order phase transition precisely accounts for the population of aggregates in mammalian cells. **E-F**: Further test of agreement between theory and super-resolution data is performed by treating cell with azetidine-2-carboxylic acid (AZC), and data from AZC treated cells in grey dots are compared directly with untreated cells (from **B & D**) represent with open circles. **E**: Upon AZC treatment, again, the data (grey) fit well to the predicted theoretical form (black line) **F**: With subtraction of surface term-fit for low n (here n<200), the resultant also remains linear in AZC treated cells suggesting that even upon perturbation the distribution changes in a manner still consistent with the physical formalism. In addition, compared to data from untreated cells (open circles),AZC data has an increased bulk (linear) term consistent with the notion that the treatment increase the degree of *super-saturation* without changing the transition mechnanism. Untreated cell data (in **B** & **D**) are from the normalized histogram of 10,000 aggregates from 10 cells and AZC data (in **E** & **F**) from 4000 aggregates from 7 cells.

The agreement between theory and experiment in Fig. 3B involves a fit with two model-parameters (surface and bulk terms respectively). We test even further whether the two terms can be decoupled. That is, whether a fit of the only the surface term at a physically appropriate limit, would result in a data-set which is accounted for primarily by the remaining bulk term.

The surface term (*an*^2/3^) must dominate for very low-n. Thus we posit that by fitting only the first few data points of the −*Log*(*P*(*n*)) graph to the surface term *an*^2/3^, and subtracting it off of the data (*25*), then for all remaining sub-critical aggregates the resultant should be the volume term,i.e. 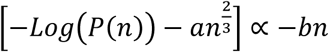 This resultant should be a straight line when plotted versus aggregate size *n*, (see theoretical prediction in Fig. 3C). We note that there is, a-priori, nothing else in our dataset imposing that the resulting data should be linear upon correction of the surface term. Thus if the data deviate from the first order phase transition energetics, we would expect a scattered resultant, or the revelation of a different energy dependence. In Fig. 3D the data show a strikingly linear resultant, demonstrating a high quality agreement between the theory and super-resolution experimental data.

From the results in Fig. 3, therefore, we conclude that while other bio-regulatory processes might be at play, the simple condensation picture with specific energetic dependence *U* (*n*) = *an*^2/3^ − *bn*, describes how sub-diffractive aggregates can form and grow to a well-defined critical size *n_c_* in the mammalian cells. We also tested aggregates with the Neuro2A cell line (neuronal precursor cells, see **supplementary text 3**) and observe the very same conclusions (**Fig. S3**) suggesting the physical mechanism for sub-diffractive aggregation may be general to a range of mammalian cells.

Biochemically, the specific parameters in the energetics for nucleation and growth would depend on the concentration of aggregating proteins and on their effective energy of interactions. To further test this notion, we sought to increase the concentration of misfolded aggregating polypeptides in living cells by incubation with a proline analog azetidine-2-carboxylic acid (AZC). This molecule incorporates in newly synthesized polypeptides instead of proline, and prevents normal folding, thus generating a massive build-up of misfolded proteins in the cell (*32*). In a condensation model, such a build-up would result in a greater degree of *super-saturation* with a stronger bulk (linear) term.

We find in Fig. 3E&F that the distribution of aggregates sizes in the presence of AZC fits the same functional form, and with a larger linear slope indicative of a larger bulk term (also see further AZC characterization in **supplementary text 4**, **Fig. S4** and general applicability of our results under other perturbations in **supplementary text 5**, **Fig. S5**). The AZC incubation data further validates the agreement between cellular aggregation data and the nucleation model, and with interpretations consistent with that expected of a classical *super-saturated* system.

Implicit to phase transition theory is the notion that any cell including healthy cells or those untreated with proteasome inhibitor, may readily form sub-diffractive aggregates which spontaneously grow into large inclusions after reaching *n_c_*. This expectation is counter to a widely held belief that the presence of precursor aggregates may directly indicate cell pathology (*2*). In Fig. 4A (left panel) we find that untreated cells do in fact show sub-diffractive aggregates implying that aggregates readily formed inside the cell without chemical treatments.

**Figure 4:**
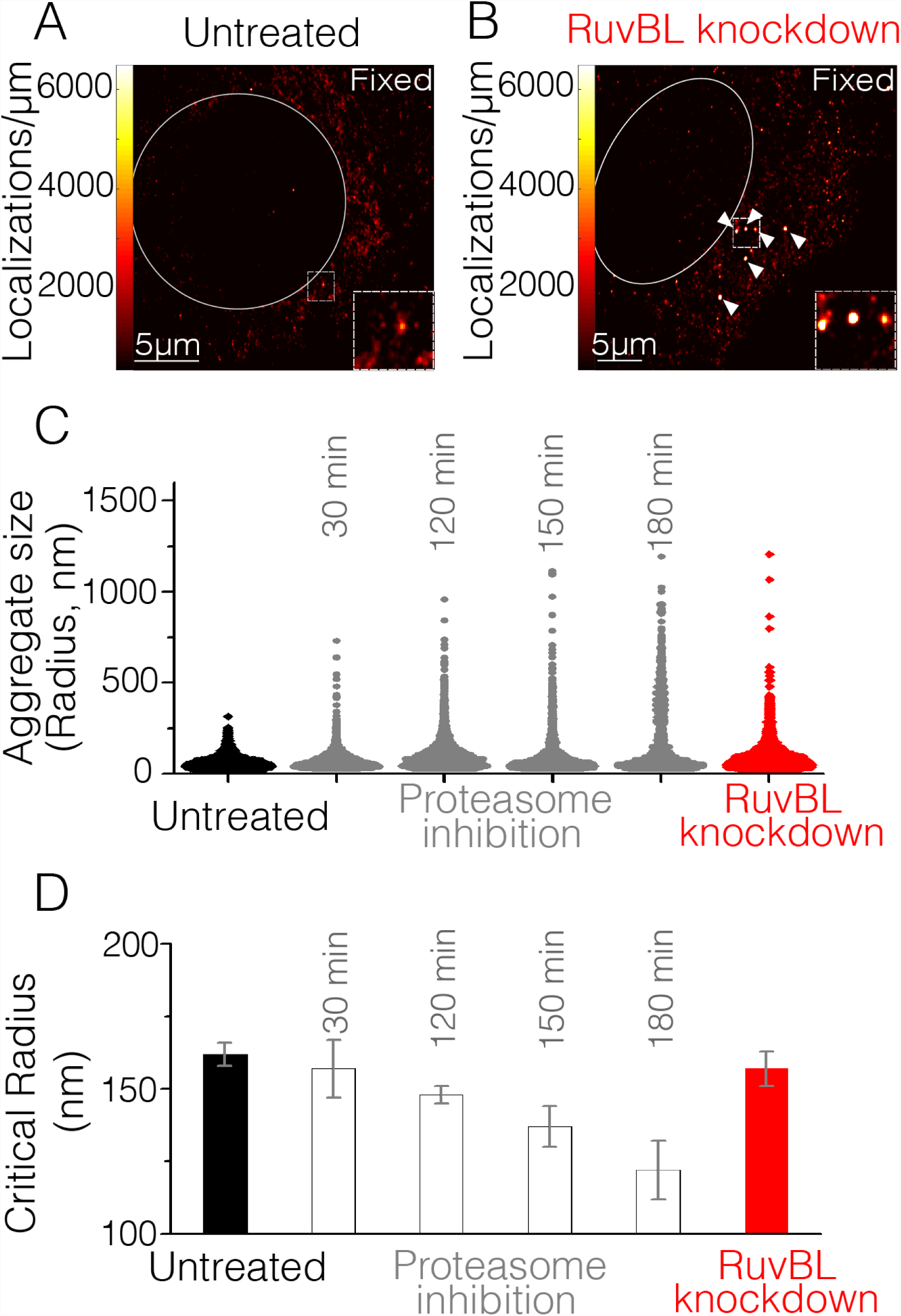
RuvBL1 dependent mechanism may clear super-critical aggregate in untreated cells without affecting the nucleation barrier. **A** Representative super-resolution reconstruction for an untreated cell visually shows many sub-diffractive aggregates (dark red) but few large aggregates (red hot). Red-hot color code is used to indicate the relative density of detections, and a white line delineates nucleus. **B** Super-resolution reconstruction from a representative cell after RuvBL1 knockdown show the cytoplasmic accumulation of relatively more intense aggregates (red hot); white arrows are used to indicate example super-critical aggregates; a red hot color code is used to indicate the relative density of detections and a white line delineates nucleus. Insets in **A, B** show zoomed in view of the largest, more intense aggregates found in each condition. **C** Violin plots showing the distribution of observed aggregate sizes (as measured by the radius) in the untreated (black), proteasome inhibited (grey) and RuvBL1 knocked-down (red) cells. Untreated cells (black) show a depletion of large clusters, and RuvBL knockdown (red) rescues the population of large clusters. **D** The critical radius (R_c_) is plotted for untreated cells (black bar), as a function of time after MG132 treatment (white bars), or upon RuvBL knockdown (red bar). RuvBL knockdown (red) did not significantly change R_c_ as compared to untreated (black) suggesting that RuvBL acted on supercritical clusters (red in **C**) without significantly changing sub-critical cluster distribution (nor the nucleation barrier). This is in contra MG132 treatments which gradually increased the population of large aggregates (gray plots in **C**) while gradually decreasing R_c_. Error bars in **D** represent errors in fit estimation as described in (*25*). Untreated cell data are from the normalized histogram of 10,000 aggregates from 10 cells, RuvBL1 knockdown from 8000 aggregates from 9 cells, and inhibition data from 5000-8000 aggregates from 6-10 cells per time point.

However, untreated cells are distinctly void of large super-critical aggregates. A violin plot of aggregate sizes from untreated cells (Fig. 4C, black) indicates that while a large population of small aggregate sizes is apparent (indicated by the width of the violin plot in Fig. 4C), untreated cells do not have a significant population of large aggregates. For instance we rarely found aggregates of radius greater than 250nm in untreated cells. We measure the critical radius to be R_c_ =162±4 nm for untreated cell (Fig. 4D, black bar, see Supplementary text 2 for calculation of *n_c_* or *R_c_*). Therefore, despite the fact that clusters should reach *n_c_* and then stably grow, such a population of super-critical clusters seems to be suppressed in healthy cells. These results imply that a hidden mechanism may exist to clear the cells of super-critical aggregates (i.e. aggregates that have reached sizes greater that the critical radius) in untreated cells.

We sought to test whether a clearance pathway could account for the absence of super-critical aggregates in untreated cells. Because a AAA+ ATPase, RuvBL, was previously suggested as a potential protein disaggregase in mammalian cells and in yeast (*33*), we tested whether RuvBL may be involved in the preferential clearance of super-critical aggregates. We find that knocking down RuvBL1 in untreated cells, results in the appearance of large aggregates (Fig. 4B, compare to untreated cell Fig. 4A). A violin plot of aggregate sizes from RuvBL1 knocked-down cells shows a clear population of large aggregate sizes, with some aggregates with radii greater than 1*μm*, a size range that could only be observed previously after hours of proteasome inhibition (Fig. 4C). These results implicate RuvBL1 in the clearance of large aggregates from untreated cells (see **supplementary text 6**, and **Fig. S6** for further tests of RuvBL1).

Importantly, we find that upon RuvBL1 knockdown, *R_c_* = 157 ± 6 *nm* did not change significantly from *R_c_* in control untreated cells (162 ±4 *nm*) (Fig. 4D) suggesting that RuvBL1 knockdown did not significantly change the sub-critical distribution. This observation implies that RuvbL1 did not affect the concentration of aggregating molecules or their interactions, unlike, for instance, proteasome inhibition which gradually reduced *R_c_* (Fig. 4D). Indeed, depletion of RuvBL1 prevented clearance of large aggregates following washout of MG132 without affecting either the distribution of aggregates in the sub-critical range or R_c_ (**supplementary text 6** and **Fig. S6**). Our data indicate that RuvBL1 dependent clearance of aggregates acts specifically on aggregates that have reached a size above *R_c_*, without changing the nucleation process.

The measured critical sizes, *R_c_*, range from ~160 nm in untreated or siRNA knockdown cells, to ~120 nm 3 hours after proteasome inhibition. These small magnitudes for *R_c_* indicate that a super-resolution technique is needed to unveil and measure this transition point *in vivo*, as by the time individual aggregates are sufficiently large to be detected in conventional cell imaging techniques they are already in the post-nucleation regime.

Previous studies bypass the direct observation of a nuclear barrier, and observe instead a sigmoidal response in the number of visible aggregates (*29*). This sigmoidal response is characterized by a lag-time followed by rapid growth after nucleation when a sufficiently large number of aggregates have crossed the nucleation barrier. However in the living cell, biological mechanisms may intervene in the post-nucleation regime to regulate the presence of larger aggregates.

Our results indicate that a hidden pathway may exist to clear cells of aggregates above the critical size, and we have identified RuvBL1 as a necessary effector in this putative super-critical clearance pathway. The mechanism by which RuvBL1, and perhaps other effectors work to preferentially clear super-critical aggregates in the cell remains currently unknown. Nonetheless, the agreement between our cellular super-resolution data and condensation systems with first order phase transition opens an avenue in the study of protein aggregation, whereby detailed theoretical predictions may be proposed and falsified experimentally, directly with quantitative *in vivo* imaging.

While our investigation has focused on aggregates related to Parkinsons disease, we note that the methodology can be readily extended to any protein that can be fluorescently tagged (for example fused to the GFP-like Dendra2). For instance, our technique may be applicable in the search for intracellular manifestations of the valency driven phase separations being studied in various contexts (*34*). Alternatively our methods could yield a thermodynamic framework underlying phase separation suggested in the formation and maintenance of membraneless organelles (*35*). Finally There is increasing evidence that the thermodynamics of first order transitions elucidated here may constitute a general organizing principle for cell biology applicable for example in cellular aging (*36*), common stress responses (*37*), and even in transcription (*38*) and other functioning of healthy cells (*39*). Thus we anticipate that this approach can help address protein aggregation directly in cells with high quantitative details, for a broad range of cellular processes and disease pathologies.

## Acknowledgments

We thank Kabir Ramola (Brandeis), Jeff Gore (MIT), Kandice Tanner (NCI/NIH) as well as members of the Cissé lab at MIT (Jan Hendrik Spille, and Micca Hecht) for helpful comments and discussions. Research reported in this publication was supported by the National Cancer Institute and the National Institutes of Health through the NIH Director’s New Innovator Award Number DP2CA195769 to IIC. The content is solely the responsibility of the authors and does not necessarily represent the official views of the National Institutes of Health. This work was also supported by funds from the MIT Department of Physics.

## Supplementary Materials

Materials and Methods

Supplementary text 1-6

Tables S1-S6

Movies S1

References (*11, 31, 30, 40-48*)

